# The hexose transporter SWEET5 confers galactose sensitivity to Arabidopsis pollen germination via the galactokinase GALK

**DOI:** 10.1101/2021.04.07.438853

**Authors:** Jiang Wang, Ya-Chi Yu, Ye Li, Li-Qing Chen

## Abstract

Galactose is an abundant and essential sugar used for the biosynthesis of many macromolecules in different organisms, including plants. Galactose metabolism is tightly and finely controlled since excess galactose and derivatives are inhibitory. In Arabidopsis, root growth and pollen germination were strongly inhibited by excess galactose. However, the mechanism of galactose induced inhibition during pollen germination remains obscure. In this study, we characterized a plasma-membrane localized transporter, AtSWEET5, that transports glucose and galactose. SWEET5 protein level started to accumulate at the tricellular stage of pollen development and peaked in mature pollen before rapidly declining after pollen was germinated. SWEET5 levels are responsible for the dosage-dependent sensitivity of galactose and galactokinase (GALK) is essential for the inhibitory effects of galactose during pollen germination. Overall, SWEET5 and GALK contribute to the maintenance of galactose metabolic homeostasis during pollen germination. SWEET5 serves as a major low-affinity hexose transporter at the early stage of pollen germination.

**One-sentence summary:** SWEET5 mediates pollen galactose sensitivity via GALK that is required for efficient galactose uptake in pollen during pollen germination.

## Introduction

Carbon flux, which is dependent on carbon supply and demand, needs to be well-controlled for plant growth and development (Hofmeyr and Cornish-Bowden, 2000; Bezrutczyk et al., 2018). Plants have developed sophisticated mechanisms to sense the sugar levels and adjust these in response to developmental or environmental cues (Ruan, 2014). Plant sensitivity to various concentrations of sugar varies depending on the specific sugar involved. For example, in *Arabidopsis thaliana*, a low 10 mM concentration of galactose can trigger an inhibitory effect on Col-0 root growth (Yamada et al., 2011; Egert et al., 2012), while the much higher 167 mM (presented as 3%) concentration of glucose or 100 mM sucrose still promotes root growth (Kircher and Schopfer, 2012; Singh et al., 2014). Additionally, during *in vitro* pollen germination, 60 mM galactose inhibits pollen germination (Hirsche et al., 2017), while 100 mM glucose does not affect the pollen germination rate (Rottmann et al., 2018) and high levels of sucrose (250-578 mM) are required for successful pollen germination (Wang et al., 2008; Rottmann et al., 2018). However, the mechanism by which galactose inhibits plant growth has not been well-characterized. Sugar transporters and enzymes are key components in the supply-demand system to maintain carbon flux control (Ruan, 2014; Julius et al., 2017). The mutants of the two hexose transporter STPs (Sugar Transport Protein), AtSTP1 and AtSTP13, have shown a galactose-tolerant phenotype for root growth (Sherson et al., 2000, Yamada et al., 2011). The transporter that is involved in the process of galactose-suppressed pollen germination, however, remains unexplored.

STPs are important during early stage gametophyte development (Truernit et al. 1999) and pollen tube growth but not in the initial phase of pollen germination (Scholz-Starke et al. 2003, Buttner, 2007). Facilitator SWEETs (Sugar Will Eventually be Exported Transporter) have been shown to transport different sugars including galactose. OsSWEET5 (Zhou et al., 2014) and CsSWEET7a (Li et al., 2021) have been demonstrated to transport galactose. So far, all plant SWEETs with a measured Km, e.g. AtSWEET1 (Chen *et al*., 2010), AtSWEET12 (Chen *et al*.,2012), AtSWEET17 (Guo *et al*., 2014), SlSWEET1a (Ho *et al*., 2019) and CsSWEET7a (Li et al., 2021), have been found to function as low-affinity glucose or sucrose transporters, ranging from ~10 to ~120 mM. It is not unusual to observe that a SWEET transports different sugars but with different efficiency and Kms (Chen *et al*., 2010, 2012, Kuanyshev et al., 2021). For example, SWEET7 has a Km of ~74 mM to glucose but ~308 mM to xylose (Kuanyshev et al., 2021). AtSWEET5/VEX, belonging to SWEET family Clade II that primarily transports hexose (Chen et al., 2010; Eom et al., 2015), is strongly expressed in the pollen vegetative cell (Engel et al., 2005), but no other details have been reported. We hypothesized that SWEET5 might be responsible for the galactose-suppressed pollen germination.

Once galactose enters a cell, it is activated by galactokinase (GALK), a cytosolic enzyme, which phosphorylates α-D-Gal into α-D-Gal-1-P at the C-1 position (Cardini and Leloir, 1953). Gal-1-P can be further converted into UDP-Gal through a reversible reaction catalyzed either by Gal-1-P uridylyltransferase (GALT) in the presence of UDP-Glc (mainly used in non-plant species) (Leloir, 1951) or by UDP-sugar pyrophosphorylase (USP) using UTP as a substrate (mainly found in plants) (Feusi et al., 1999; Kotake et al., 2004). The activated UDP-Gal can be converted reversibly into UDP-Glc by UDP-Glc epimerase (GALE/UGE) (Maxwell et al., 1960; Seifert et al., 2002). Only a single copy of the GALK gene was reported to exist in Arabidopsis (Sherson et al., 1999; Yang et al., 2009; Egert et al., 2012), and a galactose-insensitive phenotype in root growth was observed in *galk* (Egert et al., 2012). As GALK is the first enzyme to catalyze galactose, we analyzed its role together with SWEET5 transporter during pollen germination.

We show the plasma membrane localized SWEET5 transports both glucose and galactose. Data from the current study indicate SWEET5 and GALK play critical roles in controlling galactose response during pollen germination. SWEET5 transports galactose that can be phosphorylated by GALK, resulting in pollen germination inhibition through yet unknown mechanisms.

## Results

### SWEET5 transports galactose

To determine which SWEET in the mature pollen stage is able to transport galactose, we surveyed the expression and protein accumulation of all SWEETs in Arabidopsis (Mergner et al., 2020). Among all 17 members of the SWEET family, several were found to be expressed in mature pollen, but only SWEET5, SWEET11 and SWEET12 proteins were detectable there (Supplemental Figure S1). SWEET11 and SWEET12 have been reported to be transporters of sucrose (Chen et al., 2012), while SWEET5 is the most likely transporter of galactose, since its homolog OsSWEET5 has been found to transport galactose (Zhou et al., 2014). To test whether SWEET5 transports galactose, we conducted growth assays with a yeast strain EBY.VW4000 lacking 17 hexose transporters (Wieczorke et al., 1999). Yeast cells expressing SWEET5 grew on the medium containing glucose and galactose, but not fructose (Figure 1A), with slower growth on galactose than on glucose. To test whether SWEET5 transports galactose less efficiently than glucose, we conducted radio-tracer uptake, which showed that the uptake rate of galactose is lower than that of glucose (Fig. 1B). To further confirm glucose is the favored substrate than galactose to SWEET5, we conducted sugar competition experiment. Even though ten times higher concentration of galactose was used as a competitor, it was insufficient to compete with glucose uptake. Similarly, sorbitol also failed to compete with glucose uptake significantly. By contrast, the same concentration of glucose significantly reduced glucose uptake (Fig. 1C). These results support that galactose is not efficiently transported by SWEET5 relative to glucose.

**Figure 1.**
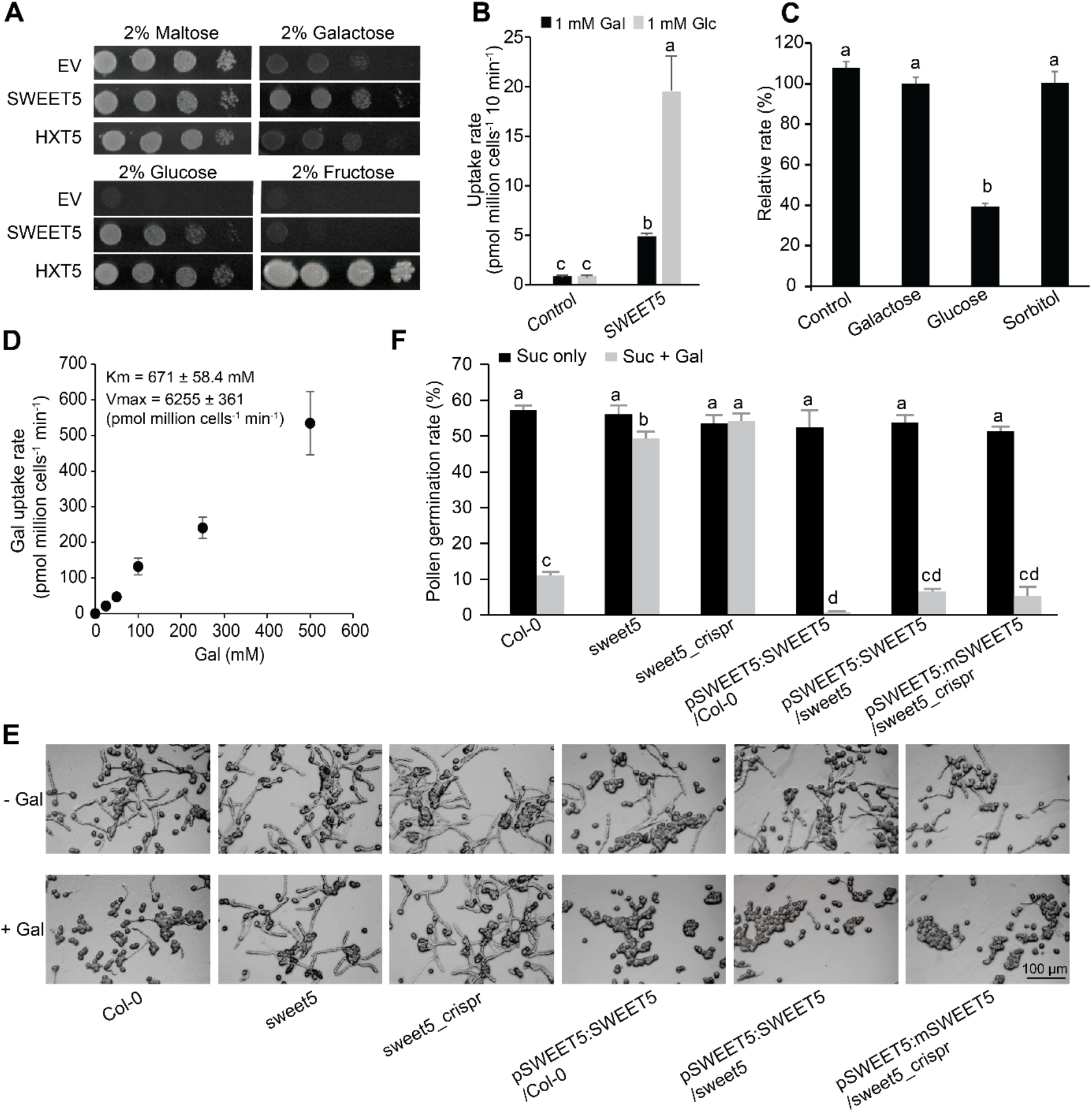
Characterization of Arabidopsis SWEET5. (A) Functional analysis of SWEET5 by EBY.VW4000 yeast complementation assay. The empty vector (EV) as a negative control, or HXT5 as a positive control. (B) SWEET5-mediated uptake of 1 mM ^14^C-galactose and ^14^C-glucose over 10 minutes in yeast cells. Cells transformed with an empty vector were used as a negative control (mean ± SE, n = 4). (C) Substrate competition uptake analysis of SWEET5 in yeast over 10 min. Different 50 mM cold sugars competed with 5 mM cold glucose (including ^14^C-glucose) that was normalized to 100 % (mean ± SE, n = 4). (D) Kinetics of ^14^C-galactose accumulation by SWEET5 in yeast (mean ± SE, n = 4). An empty vector served as a negative control and was subtracted. (E) *In vitro* pollen germination assay for various genotypes on sucrose-based medium with or without 60 mM galactose. The pictures were taken 8 hours post-germination. (F) Statistical analysis of pollen germination rates for various genotypes. The means were calculated from multiple repeats (± SE, n ≥ 9), with over 450 pollen grains/tubes counted in total. The statistically significant differences among different samples were determined using one-way ANOVA followed by multiple comparison tests and were represented by different letters (*P* <0.05).

Additionally, we did the kinetic analysis of SWEET5. The Km for galactose uptake of SWEET5 was calculated as 671 ± 58.4 mM (Figure 1D), however, we do not know whether osmolarity will affect the physiological performance of yeast cells when cultured with a high concentration (>500 mM). Therefore, to test if the osmolarity will affect yeast uptake and thus calculated Km, we compared galactose uptake with and without osmolarity adjusted by mixing different designed concentrations of galactose with matched concentrations of sorbitol to reach a total concentration of 500 mM and observed no significant osmotic effects on galactose uptake. (Supplemental Figure S2).

### SWEET5 contributes to galactose inhibition of Arabidopsis pollen germination

To test how galactose affects pollen germination under our setting, we performed pollen germination experiments using pollen germination medium (PGM) supplemented with 60 mM galactose compared with that supplemented with or without other sugars. The pollen germination rate of Col-0 was dramatically reduced from 57% to 11% when PGM with 60 mM Galactose (Figure 1E and 1F), while remained unchanged when PGM was supplemented with either 60 mM glucose or fructose (Supplemental Figure S3). The galactose suppression was barely detectable in *SWEET5* loss-of-function mutants created from T-DNA insertion (*sweet5*) or CRISPR/CAS9 editing (Figure 1E and 1F; Supplemental Figure S4). The phenotypes of the mutants were fully complemented by transformation with a construct carrying the *SWEET5* promoter driving either endogenous *SWEET5* or a synonymous m*SWEET5* that carries mutations to avoid gRNA recognition in CRISPR mutant lines (p*SWEET5*:*mSWEET5/sweet5_crispr*; Figure 1E and 1F; Supplemental Figure S4D). Accordingly, the SWEET5 overexpression lines (p*SWEET5*:*SWEET5*/Col-0) were more sensitive to galactose than Col-0 (Figure 1E and 1F). To test whether in vivo pollen germination is affected in *sweet5* mutant, we compared pollen germination and pollen tube growth of *sweet5* with that of Col-0 on Col-0 pistil after hand-pollination. No obvious differences in the pollen germination/pollen tube growth (aided by aniline blue staining) in vivo were observed at 2 hours or 6 hours after pollination (Supplemental Figure S4G), suggesting the galactose content at the apoplasm of stigma cells is not high enough to trigger pollen inhibition under normal condition. It is worth noting that glucose has a marginal effect on pollen germination (Supplemental Figure S3) and thus is negligible, even if present. However, it is not known if galactose content will rise to affect pollen germination under stress conditions. For example, soluble galactose was significantly increased in coffee plant leaves under heat stress, likely due to cell-wall modifications (Lima et al., 2013). Moreover, there were no observable morphological phenotypes in *sweet5* mutant under normal conditions, which may be due to functional redundancy with other plasma membrane-localized hexose transporters (STPs, PMTs) detected at either the RNA or protein level in pollen (Supplemental Figure S5). Among those hexose transporters that were detected at the protein level, many can transport glucose and galactose (Supplemental Table S1).

### SWEET5 is mainly expressed at the late stages of pollen development

To determine the timing of the initiation of SWEET5-associated galactose inhibition, we examined the spatial and temporal accumulation patterns of SWEET5 using transgenic lines harboring a SWEET5 translational fusion with β-glucuronidase (GUS) or yellow fluorescent protein (YFP) driven by its native SWEET5 promoter (p*SWEET5*:*gSWEET5-GUS* or p*SWEET5*:*gSWEET5-YFP*). SWEET5 was almost exclusively found in the anthers and mature pollen at late stages of flower development (Figure 2A), although GUS staining was detected at the early seedling stage with a preference in the vein. The SWEET5 protein accumulation pattern was carefully assessed in the flowers. Flowers at different developmental stages (Bowman, 1994) were imaged using fluorescence microscopy to detect SWEET5 protein levels using the reporter YFP. As shown in Figure 2C, SWEET5 started to accumulate in immature pollen grains from flower stage 11, in which pollen mitosis I and II occur (Cecchetti et al., 2008) and peaked at flower stage 13 (anthesis). The YFP signal was also observed over the course of male gametophyte development by isolating developing pollen grains. The SWEET5-YFP protein was first detected at the tricellular pollen (TCP) stage and peaked at the mature pollen grain (MPG) stage, while it was undetectable in both uninucleate microspores (UNM) and bicellular pollen (BCP), as delineated by PI staining of the cell wall and nucleus (Figure 2B). The observed pattern of SWEET5-YFP protein agrees with the reported gene expression profiling of *SWEET5* in Arabidopsis pollen over different developmental stages (Honys and Twell, 2004), namely that *SWEET5* was highly expressed at the tricellular stage and mature pollen stage. Additionally, SWEET5 protein level was substantially reduced in germinated pollen/pollen tube (marked by white arrow) compared with non-germinated pollen grains after *in vitro* pollen germination (Figure 2D). The subcellular localization of SWEET5 was determined using pollen from transgenic plants harboring a SWEET5 (CDS) translational fusion with a green fluorescent protein (GFP) driven by its native SWEET5 promoter (p*SWEET5*:*cSWEET5-GFP*). SWEET5 is localized to the plasma membrane, and also endomembrane compartments as delineated by Nile red staining for the lipid droplets (Figure 2E). It is not unusual to observe that strong fluorescence throughout the cell, including endomembrane, when a tagged protein is abundant in pollen (Tunc-Ozdemir et al., 2013). This has been similarly observed for other plasma membrane localized proteins in pollen (Frietsch et al., 2007; Tunc-Ozdemir et al., 2013; Hamilton et al., 2015). SWEET5 localization was further transiently examined using the reporter mVenus in *N. benthamiana* leaves (Gookin and Assmann, 2014). SWEET5 protein was localized to the pavement cell periphery enclosing both chloroplast and Golgi apparatus (Figure 2F), which suggests SWEET5 localized to the plasma membrane, consistent with published Arabidopsis SWEETs except for SWEET2, SWEET16, and SWEET17 that are localized to the tonoplast (Klemens et al., 2013; Guo et al., 2014; Chen et al., 2015a).

**Figure 2.**
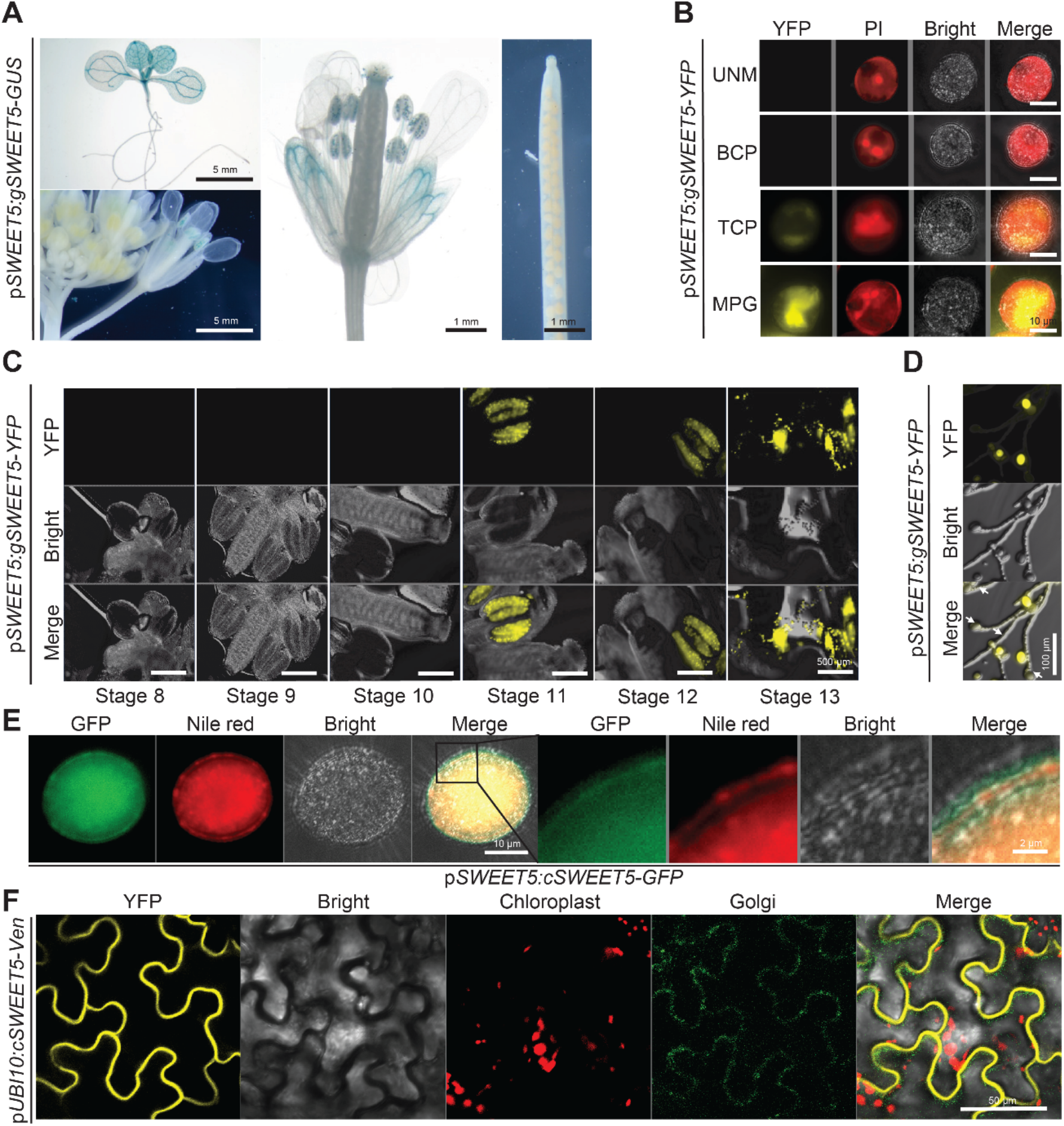
Tissue-specific accumulation and cellular localization of SWEET5. (A) SWEET5 tissue-specific expression was evaluated. Twelve-day-old seedlings, inflorescences and siliques from about one-month old Arabidopsis carrying p*SWEET5*:*gSWEET5-GUS* were histochemically stained 4 h for GUS activity. (B) SWEET5 localization in pollen grain was examined using fluorescence microscopy. Images were captured using the YFP filter for YFP, the RFP filter for propidium iodine (PI) staining, and under bright field. SWEET5 was detected in the tricellular pollen grain (TCP) and mature pollen grain (MPG) but absent in early stages of uninucleate microspores (UNM) and bicellular pollen (BCP). (C) SWEET5 accumulation in different floral stages of Arabidopsis was examined using fluorescence microscopy. The signals started to be detected from stage 11. (D) SWEET5 accumulation was detected after pollen germination in p*SWEET5*:*gSWEET5-YFP lines*. Pictures were taken 8 hours post-germination using a fluorescence microscope. White arrows pointed to germinated pollen with much weaker fluorescence than un-germinated pollen. (E) SWEET5 cellular localization was examined in mature pollen carrying p*SWEET5*:*cSWEET5-GFP*. (F) SWEET5 cellular localization was examined 72 hours after infiltration of *Nicotiana benthamiana* leaves with *pUBI10*:*cSWEET5-mVenus*. Pictures were taken using confocal microscopy.

### Galactose sensitivity is GALK-dependent during pollen germination

To further elucidate whether GALK is involved in the galactose sensitivity during pollen germination, we first surveyed whether GALK is accumulated in the pollen. The RNA transcripts of *GALK* are abundant in all pollen developmental stages, preferentially in microspore and bicellular pollen stages (Honys and Twell, 2004). The protein level of GALK is relatively consistent across all tissues, including mature pollen grains (Supplemental Figure S6A). Therefore, we tested pollen germination of the loss-of-function *galk* mutant (Egert et al., 2012) on PGM containing galactose. The *galk* mutant showed a galactose-tolerant phenotype similar to that of *sweet5* (Figure 3A; Supplemental Figure S7A), although it is a knock-down mutant with *an* 80% reduction in mature pollen RNA transcripts (Supplemental Figure S6C). The *galk* galactose-tolerant phenotype was fully complemented by *GALK* expressed under control of the pollen-specific *LAT52* promoter (Figure 3A; Supplemental Figure S7A). Notably, the *galk* complementation line showed a much stronger galactose sensitivity than Col-0, which is likely due to the over-expression of GALK in the complementation line (Supplemental Figure S6C). As expected, the GALK and SWEET5 double mutant *sweet5 × galk* showed a similar galactose-tolerant phenotype to their individual single mutants, which indicates that GALK and SWEET5 function in the same pathway way. As Gal-1-P is suggested to exert cellular toxicity from many studies in animals (Lai et al., 2009), it is reasonable to speculate that pollen sensitivity to galactose may be due to the accumulation of potential cytotoxic Gal-1-P, which depends on GALK activity (de Jongh et al., 2008). To test this possibility, we compared the effects Gal-1-P on pollen germination, 6 mM Gal-1-P was sufficient to mimic the suppression effect produced by 60 mM galactose in Col-0 (Figure 3B; Supplemental Figure S7B). Gal-1-P can be converted into UDP-Gal by USP in plants (Feusi et al., 1999). To test whether the USP-catalyzed reaction could alleviate galactose suppression, we generated over-expression lines of AtUSP driven by the pollen-specific *pLAT52* promoter in Col-0 background. However, USP overexpression lines failed to improve the pollen germination rate under 60 mM galactose condition (Supplemental Figure S8).

**Figure 3.**
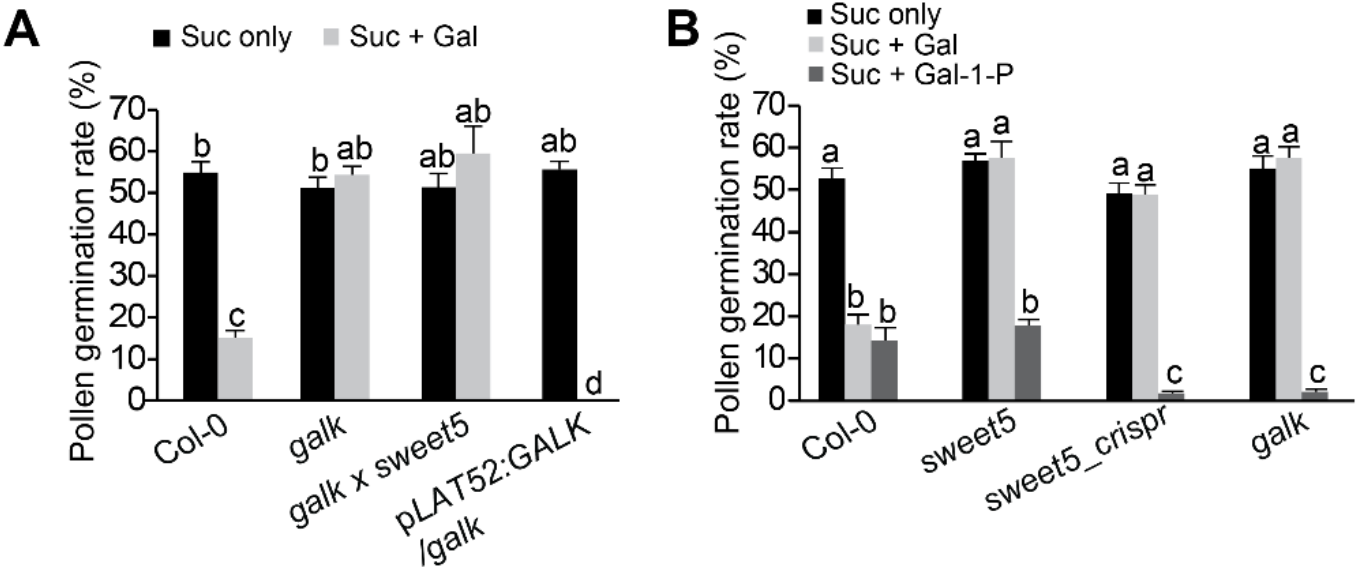
Involvement of GALK in galactose-inhibited pollen germination (A) Examination of *in vitro* pollen germination on sucrose-based medium with 0 mM or 60 mM galactose for various genotypes. (means ± SE, n ≥ 6 (over 300 pollen grains/tubes counted) and *P*<0.05). (B) Effects of galactose-1-phosphate (6 mM) on *in vitro* pollen germination (means ± SE, n ≥ 6 (over 300 pollen grain/tubes counted) and *P* <0.05).

### Galactose inhibition is dosage-dependent

To determine the range of concentration at which galactose inhibits pollen germination, pollen from different genotypes were germinated on PGM with galactose concentrations ranging from 0 to 600 mM. In Col-0, the pollen germination rate was not affected under 0.6 mM galactose, started to decline significantly at 6 mM, followed by a substantial decrease at 60 mM and further reduced to almost zero at 600 mM (Figure 4A, Supplemental Figure S7C). A time-course pollen germination experiment of Col-0 further confirmed that the pollen germination rate was significantly affected in the presence of 6 mM galactose starting from four hours after germination (Supplemental Figure S7D). However, no significant difference was observed between Col-0 and *sweet5* mutants at 6 mM (Figure 4A, Supplemental Figure S7C). *sweet5* mutants were generally tolerant to galactose up to 60 mM. Consistent with the observation, the SWEET5 overexpression lines were more sensitive to ≥ 6 mM galactose than Col-0, with pollen germination rates reduced to near zero by 60 mM galactose. These results suggest that SWEET5 is responsible for galactose flux at the higher concentration range. To assess the differences in galactose uptake between Col-0 and *sweet5*, we conducted ^14^C-galactose tracer uptake assays in pollen with various galactose concentrations. Pollen grains germinated in a liquid PGM for 45 mins (the hydration period, Wang *et al*., 2008), minimizing the interference from transporters expressed on pollen tubes. Less ^14^C-galactose was taken up by *sweet5* pollen than by Col-0 pollen at 60 mM galactose concentration (Figure 4B). By contrast, differences were not prominent at lower amounts of galactose (Figure 4B).

**Figure 4.**
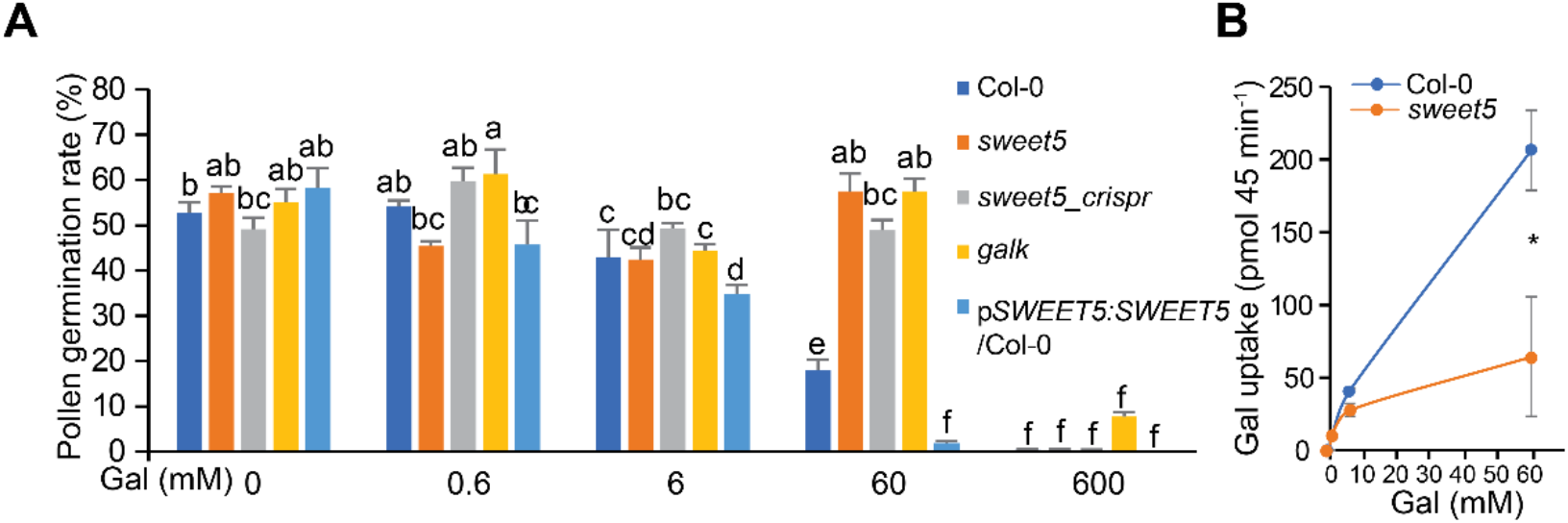
Concentration-dependent galactose response. (A) *In vitro* pollen germination of various genotypes on medium supplemented with different concentrations of galactose (ranging from 0 to 600 mM). The means (± SE) of at least 6 repeats were plotted (over 300 pollen grains/tubes tested) (*P* < 0.05). (B) Concentration-dependent galactose uptake in pollen of Col-0 and *sweet5* (means ± SE, n=3 and *P* < 0.05). Pollen germinated on liquid medium containing 0 mM, 0.6 mM, 6 mM, or 60 mM galactose in addition to 0.3 μCi ^14^C-galactose per sample for 45 min.

Galactose at 10 mM was reported to strongly inhibit root growth of Col-0 (medium containing no sucrose, Yamada et al., 2011; Egert et al., 2012), while higher galactose (60 mM) is needed to strongly inhibit Col-0 pollen germination (medium containing 15% (~ 439 mM) or 19.8 % (~578 mM) sucrose, Hirsche et al., 2017 or current study). To test whether the sucrose in the medium may differentially affect the observed galactose response, which may lead to a biased comparison, we conducted a galactose dosage-dependent assay for root growth using Col-0. Root growth was not affected at 1 mM galactose but was strongly inhibited at 6 mM or higher without sucrose in the medium (Supplemental Figure S9). By contrast, a significant root reduction was only observed at 60 mM galactose when 0.5% or 1% sucrose was supplemented in the medium. Therefore, the level of inhibitory galactose concentration should be considered in the context of sucrose presence or absence.

## Discussion

### Pollen is a simple system to study the control of galactose flux

While pollen has a fundamental role in the sexual reproduction of flowering plants, pollen germination together with pollen tube growth has served as a model system to study single cell growth and morphogenesis (Feijó et al., 2004). In Arabidopsis, pollen grains are highly homogeneous and have strikingly low transcriptional complexity (on average, 6044 genes expressed in mature pollen; Rutley and Twell, 2015) compared to sporophytic tissues or purified sporophytic cells (e.g., 13,222 genes expressed in stomatal guard cells, and 11,696 genes expressed in root hair cells; Bates et al., 2012, Becker et al., 2014). Our work to understand relationships among galactose transport, GALK, and galactose sensitivity sheds light on developing alternative strategies to attenuate galactose suppression by fine-tuning their relationships.

Over the past several decades, extensive efforts have been made to study galactose toxicity in humans (galactosemia) and yeast; however, the mechanism underlying the toxicity is far from understood (Lai et al., 2009). Considerably less attention has been paid to galactose inhibition by plant scientists. In the current study, we found that the plasma membrane-localized hexose transporter AtSWEET5 is responsible for galactose sensitivity during pollen germination. A high concentration of sucrose at 19.8% is needed in the medium for *in vitro* pollen germination, which may be partially cleaved into glucose and fructose by the Arabidopsis cell-wall invertase 2 before import into pollen (Hirsche et al., 2009). As SWEET5 prefers to transport glucose (Figure 1B), glucose may competitively suppress galactose uptake, resulting in much less galactose transported in pollen. Thus, a high concentration of galactose, such as 60 mM, is needed to result in a severe inhibitory effect on pollen germination. This concept was supported by the reduced galactose response during root growth of wild-type plant in presence of 0.5 % sucrose (Supplemental Figure S9). Even when 60 mM external galactose was used, the estimated concentration was only about 135 μM in Col-0 pollen after incubation with 60 mM galactose for 45 min. This was calculated based on the uptake of 200 pmol galactose (Figure 4B) determined from 20 flowers that yielded a combined 40,000 pollen (2000 pollen per flower), and the mean pollen volume deduced from the Arabidopsis pollen diameter of 20.72 μm (De Storme et al., 2013).

### Model of galactose transport and metabolism during *in vitro* germination of pollen

From the data presented in this study and literature reports, a schematic model of galactose sensitivity in pollen was illustrated in Figure 5: SWEET5, as a uniporter, transports galactose across pollen membrane across a concentration gradient. If galactose is not sufficiently transported into pollen grains due to a lack of enough SWEET5 protein, a galactose-tolerant phenotype can be observed. Similarly, the lack of AtSTP1 and AtSTP13 have shown a galactose-tolerant phenotype of root growth in conditions of up to 100 mM galactose (Sherson et al., 2000, Yamada et al., 2011). Once galactose is taken up by cells, it will first be phosphorylated by GALK to produce the potentially toxic Gal-1-P. A galactose detoxification pathway that stores excess galactose in the vacuoles was proposed in the *galk* mutant (Egert et al., 2012). This pathway of detoxification cannot explain the galactose-tolerant phenotype of *galk* pollen, because vacuoles are nearly absent from mature pollen grains (Pacini et al., 2011). The disrupted Gal-1-P production may therefore be responsible for the galactose tolerance of the *galk* mutant during pollen germination. Though possible free galactose accumulation by *galk* pollen, similar to *galk* leaves (Egert et al., 2012), may negatively impact galactose influx during pollen germination and obscure our understanding.

**Figure 5.**
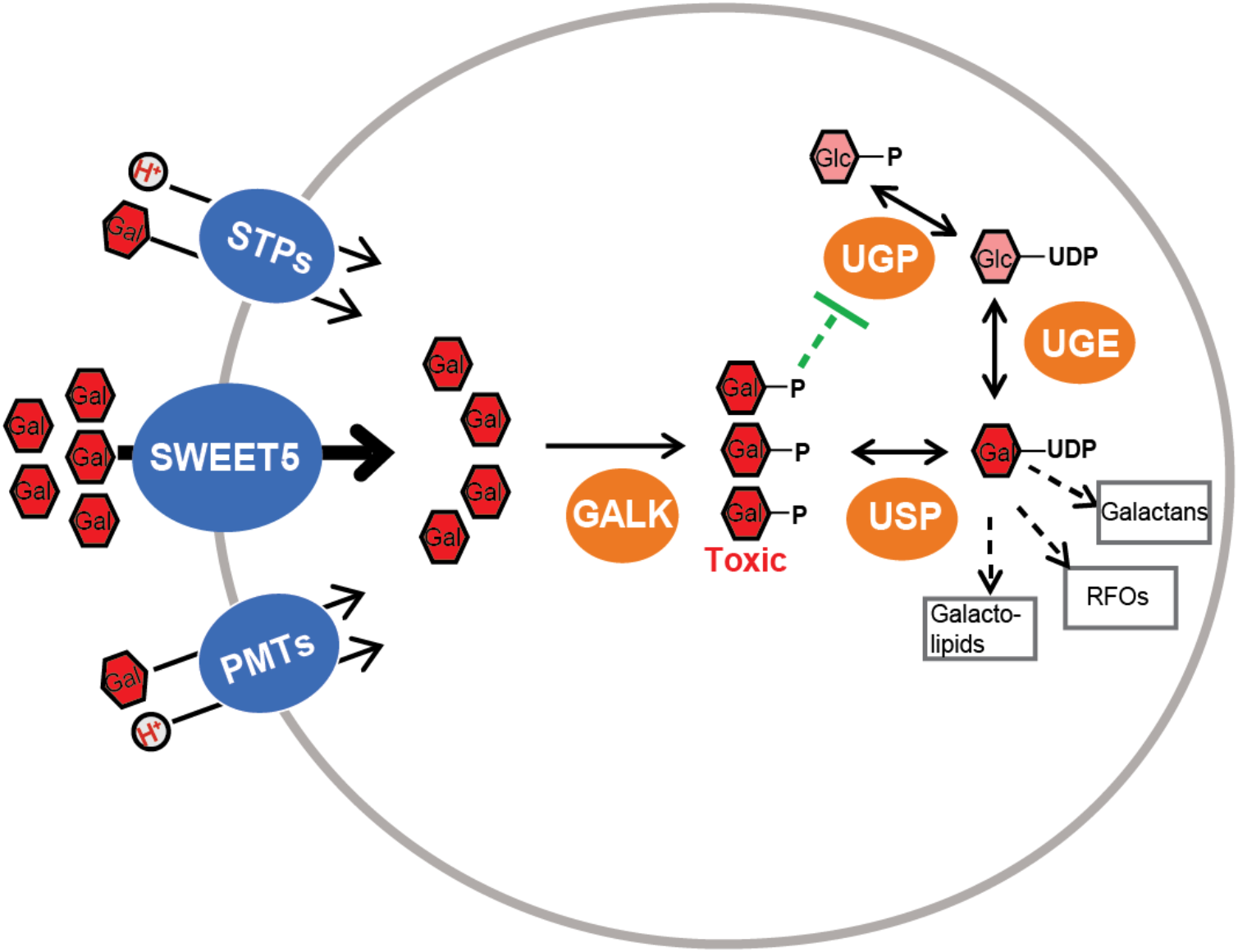
Schematic representation of galactose transport and metabolism in Arabidopsis pollen. A simplified pollen structure was drawn for clarity. Arrows represent positive regulation and bar-headed lines indicate negative regulations. The black lines represent galactose metabolic flux. The thick black line means higher flux. The green dotted line is for hypothetical regulation. Galacto-lipids are predominantly found in plastid membrane. RFOs (Raffinose Oligosaccharides) mainly include raffinose and stachyose containing one and two Galactose moieties, respectively. Galactans represents the cell wall-bound galactose.

Our study suggested that Gal-1-P is responsible for the galactose sensitivity observed during pollen germination, which is supported by the results of Gal-1-P effect on *in vitro* pollen germination. However, caution should be taken when interpreting these results, because it is still unclear how Gal-1-P can be taken up by pollen as no such transporters have been identified. Gal-1-P showed a significant increase as the external galactose concentration rises beyond the inhibitory levels in corn and barley coleoptiles (Roberts et al., 1971). In animals or yeast, Gal-1-P has been reported to potentially inhibit enzymes such as phosphoglucomutase (de Jongh et al., 2008), UDP-glucose pyrophosphorylase (UGPase, catalyzing the reversible conversion from Glc-1-P to UDP-Glc) (Lai et al., 2003), and inositol monophosphatase (Bhat, 2003) *in vitro*, however, no *in vivo* targets of Gal-1-P have been identified. In limited studies in plants, Gal-1-P has only been reported to inhibit UGPase activity in the crude extract from oat coleoptiles, but not from the azuki bean epicotyl, suggesting that galactose sensitivity appears to vary depending on the species (Yamamoto et al., 1988). We attempted to measure the galactose-associated metabolites upon galactose treatment in *sweet5* and *galk*, however, no clear conclusions can be drawn due to the difficulties of separating glucose and galactose conjugated phosphates and nucleotide sugars using an LC-MS/MS system.

### SWEET5 orthologs may play essential roles in hexose-utilizing pollen

Compared to the proton sugar symporters, SWEET5 likely functions as a uniporter, facilitating substrate movement along a substrate concentration gradient (Chen et al., 2015b). The physiological importance of SWEET family members was underscored mainly by their involvement in multiple processes requiring sugar transport capability, for instances, SWEET11 and SWEET12 in apoplasmic phloem loading (Chen *et al*., 2012); SWEET9 in nectar secretion (Lin *et al*., 2014); and SWEET11, SWEET12, and SWEET15 in seed filling (Chen *et al*., 2015c). In the current work, we showed that SWEET5 is highly expressed in the mature pollen but dramatically declines during the transition from germinated pollen to pollen tubes, suggesting that SWEET5 functions during the early stages of pollen germination. By contrast, several STPs are primarily expressed in pollen tubes but are almost absent in mature pollen grains (Rottmann et al., 2018). For example, STP10 is a pollen tube-specific hexose transporter (Rottmann et al., 2016); the glucose transporter STP9 is weakly expressed in mature pollen grains and is primarily expressed in growing pollen tubes (Schneidereit et al., 2003). Therefore, SWEET5 and STPs may have complementary roles during pollen germination. As reported, high concentrations of sugars (ranging from 2% to 19.8%) are needed for successful germination *in vitro* in various species (Ylstra et al., 1998; Stadler et al., 1999; Wang et al., 2008) since pollen germination is an energy-demanding process (Reinders, 2016). However, the effect of different sugars on *in vitro* pollen germination depends on the species. For example, sucrose and hexose differentially affect *in vitro* pollen germination, ranging from growing equally well on glucose, fructose, or sucrose for petunia (Ylstra et al., 1998), cucumber (Cheng et al., 2015), and pearl millet (Reger et al., 1992), to a nearly complete inhibition by glucose or fructose for Arabidopsis (Hirsche et al., 2017). But for some species, such as tobacco, either sucrose or glucose can support pollen germination, while only sucrose can promote pollen tube growth (Goetz et al., 2017). Galactose alone can also promote pollen germination, for example in *Picea wilsonii* (Zhou et al., 2020) and cucumber (Cheng et al., 2015) to a similar extent as sucrose. Therefore, the lack of a phenotype in *sweet5* under normal conditions may be due to the preference of Arabidopsis pollen during the initial phase of germination for sucrose. Considering SWEET5 also transports glucose, it is rational to speculate that SWEET5 orthologues may be essential for pollen development, and therefore seed production in the species that prefer to use hexose as a carbon source during pollen germination or pollen tube growth.

## Conclusions

We have presented SWEET5 as a plasma membrane-localized hexose transporter that is highly expressed in mature pollen grains. GALK catalyzes the synthesis of Gal-1-P from galactose that can be transported by SWEET5. The pollen grains of *sweet5* and *galk* mutants are tolerant to galactose inhibition during pollen germination in vitro, suggesting that SWEET5 and GALK play a critical role in the sensitivity of pollen germination to galactose in Arabidopsis. Pollen germination studies served as a tool to elucidate SWEET5 biological functions and provide a model system that facilitates research to uncover mechanisms underlying galactose sensitivity. The future work associated with either SWEET5 or its homologs from species that favor different carbon sources for pollen germination, will offer new insights into the carbon skeleton and/or energy supply mechanisms responsible for successful reproduction.

## Materials and Methods

### Constructs for yeast expression

The CDSs of *AtSWEET5* were amplified by RT–PCR using gene specific primers named by gene (Supplemental Table S2, P1 and P2) and were subsequently cloned into the donor vector pDONR221-f1 by BP clonase (Invitrogen) before transfer to pDRf1-GW (Loque et al., 2007) by LR clonase (Invitrogen).

### GUS and YFP fusion constructs under native promoters

To construct *SWEET5* fused with GUS (p*SWEET5*:*gSWEET5-GUS*), the fragment including the *SWEET5* native promoter sequence (1801 bp upstream of ATG) and genomic sequence of *SWEET5* was amplified (P3 and P4) and cloned into pDONR221-f1 by BP reactions followed by recombination into the vector pMDC163 carrying the GUS gene (Curtis and Grossniklaus, 2003) by LR reaction. For the SWEET5 overexpression construct (p*SWEET5*:*gSWEET5-YFP*), the entry clone from above was transferred into pEarleyGateTW1 (Wang et al., 2016) by LR reactions.

### eGFP fusion constructs under different promoters for strength comparisons and mutant complementation

The *SWEET5* (1216 bp upstream of ATG) native promoter, and *LAT52* promoter (602 bp; Muschietti et al., 1994) were amplified with specific forward primers (P5-P8) containing a *BamH*I restriction site and the reverse primers containing an *Xba*I restriction site and sub-cloned into the eGFP-containing vector pGTKan3 (Kasaras and Kunze, 2010) via *BamH*I and *Xba*I restriction sites. The CDS of *SWEET5, GALK*, and *USP* by their own specific primers (P9-P14) were seamlessly subcloned into corresponding pGTKan3_promoter constructs by In-Fusion® (Takara) following the linearization at *Xba*I and *Pst*I restriction sites. For the genome editing construct of *SWEET5*, gRNA that targets the first exon of SWEET5 (Supplemental Figure S4D) was cloned into p*YAO*:*hSpCas9* (Yan et al., 2015). For the synonymous complementation construct of Cas9 containing *sweet5_crispr*, synonymous *SWEET5* driven by pSWEET5 was amplified by primer P15 and P16 based on pDONR221_pSWEET5-*SWEET5* with gRNA targeting the region modified through site-directed mutagenesis before transferring into pEarleyGate301 (Earley et al., 2006) by LR reactions.

### Construct for subcellular localization

For subcellular localization of SWEET5, after the vector pDOE-13 (Gookin and Assmann, 2014) was linearized by *SanD*I + *AatI*I restriction enzymes at MCS3, the CDS of SWEET5 was amplified by primers P17 and P18 and subcloned into it by In-Fusion® (Takara). The agroinfiltration-based assays were performed as previously described (Gookin and Assmann, 2014). The *N. benthamiana* leaf disks were collected 72 h post infiltration for confocal imaging. Three independent trials were conducted.

### Plant materials and growth conditions

The Arabidopsis Col-0 plants were grown under controlled temperature (22°C) with a 16-h light (100-150 μEm^−2^s^−1^)/ 8-h dark photoperiod. The flowers at stage 13 (fully opened) were collected at around 5 hours into the light period for different experiments. T-DNA mutants of *galk* (GABI-Kat 489D10), and *sweet5* (CS853155) were obtained from NASC or ABRC. Homozygous lines were genotyped using primers of P19-P24 and used in related experiments. The floral dip method (Clough and Bent, 1998) was used to generate all the transgenic lines used in this study. At least 15 T_1_ lines were generated for each construct and at least 3 randomly selected lines were propagated to generate T_3_ homozygous seeds.

### Yeast complementation assay and ^14^C uptake assay

Yeast complementation and yeast ^14^C uptake assays were conducted following the previously described method (Chen et al., 2010). ^14^C-glucose (0.1 μCi D-[U-^14^C] glucose; 275 mCi/mmol), ^14^C-galactose (0.1 μCi D-[1-^14^C] galactose; 56.2 mCi/mmol) was added per sample and four independent transformants were used for each uptake experiment. For concentration- and time-dependent uptake of [14C] galactose for SWEET5, 5 galactose concentrations (25, 50, 100, 250, 500 mM), and 4 time points (10 sec, 2.5, 5, and 10 min) for each concentration were used to collect yeast cell samples and measure galactose uptake. Sugar uptake at 10 sec was normalized as 0.

### *In vitro* pollen germination

The *in vitro* pollen germination assay was conducted according to a previously described method (Wang et al., 2008) with minor modifications. The sucrose only basic pollen germination medium (PGM) was composed of 19.8% (w/v) Suc (15% (w/v) Suc in liquid PGM), 1.5 mM boric acid, 0.8 mM MgSO_4_, 1 mM KCl, 5 mM MES, 0.05% (w/v) lactalbumin hydrolysate, 10 μM myoinositol, 5 mM CaCl_2_ and 1.5% (w/v) agarose. The pH was adjusted to 5.8 using 1 mM Tris (pH = 8). The medium solution was heated to boiling on a heat plate, and then 500 μl was spread evenly onto 75 × 25 mm glass slides. Slides were cooled down at room temperature before placing in an opaque, sealed slide box. The medium was made freshly for each germination experiment. Six flowers at stage 13 were collected from each genotype to spread pollen gently onto a ~1 cm^2^ area of the germination medium. The slide box was kept at room temperature for 8 hours in the dark samples on slides were imaged with a compound microscope. Pollen with pollen tubes longer than the diameter of the pollen grain (about 20 μm) was considered as germinated pollen.

### Pollen galactose uptake

The liquid pollen germination method was adopted from a previously described method (Wang et al., 2008) with minor modifications for ^14^C-galactose uptake. For each trial, 20 flowers (at least 2000 pollen grains collected per flower) at stage 13 for each genotype were collected into a 2 ml tube, and then 1 ml of liquid PGM was added in and vortexed vigorously for 60 s to release mature pollen grains into the solution. Subsequently, the pollen grains were counted using a hemocytometer. Equal amounts of pollen for all tested genotypes were collected by centrifuging at 15,000 g for 1 min. The pollen pellet was resuspended in 30 μl of liquid PGM containing ^14^C-galactose (0.3 μCi hot galactose) as well as different concentrations of cold galactose (ranging from 0 to 60 mM), and subsequently cultured in a petri dish (35 mm in diameter). The petri dish was covered and placed in the dark for 45 min before pollen was washed using 1 ml of ice-cold liquid PGM and collected into a 1.5 ml tube. The pollen was precipitated at 15,000 g for 1 min and the pellet was washed by 1 ml ice-cold pollen isolation buffer (PIB, composed of 100 mM NaPO^4^, pH 7.5, 1 mM EDTA, and 0.1% (v/v) Triton X-100) 5 more times. The pellet was resuspended in 100 μl PIB and transferred to a scintillation vial containing 5 ml Ultima Gold XR Scintillation liquid (PerkinElmer). Radioactivity for each vial was measured by liquid scintillation spectrometry.

### GUS histochemical analysis

GUS staining was performed as previously described (Chen et al., 2012). Twelve-day-old seedlings were used, and inflorescences and siliques were collected for histochemical GUS staining.

### Microscopy imaging

A Zeiss Apotome.2 (Carl Zeiss, Thornwood, NY, USA) was used for fluorescence acquisition. Different FL filter sets (YFP, CFP, RFP and GFP) were used to image samples as needed. A Zeiss LSM 710 confocal microscope (Carl Zeiss, Thornwood, NY, USA) was used to image samples. Argon laser excitation wavelength and emission bandwidth were 405 nm and 425-550 nm for aniline blue, 458 nm and 480–520 nm for mTq2 (cyan), 488 nm and 500–550 nm for mVenus (yellow), and 633 nm and 633-740 nm for chlorophyll autofluorescence (red) respectively. Image acquisition parameters were held consistent. Raw data from each channel were not altered beyond equal signal increases.

### RNA isolation and RT-qPCR

RNA isolation from pollen was performed as previously described with modifications (Lu, 2011). For each independent sample, pollen grains from 50 flowers at stage 13 were collected. For samples under galactose treatment, pollen grains from 50 flowers were germinated in liquid PGM for 45 mins as described above in pollen galactose uptake. RNA was isolated using Trizol (Invitrogen) as instructed by the manufacturer. First-strand cDNA was synthesized using oligo(dT) and M-MuLV reverse transcriptase (NEB). Real-time qPCR was performed using PowerUp™ SYBR™ master mix (Applied Biosystems) according to the manufacturer’s instructions on a CFX96 Real-Time PCR Detection System (Bio-Rad) using gene specific primers (P25-P30). The expression values were normalized to *ACTIN8* expression values in each repeat, and subsequently normalized to Col-0 using 2^−ΔΔ CT^ method (Livak and Schmittgen, 2001).

### Statistical analysis

The differences between the two subjects were determined using the two-tailed Student’s t-test with equal variance. The differences among multiple subjects were assessed using one-way ANOVA followed by multiple comparison tests (Fisher’s LSD method). All statistical analysis was performed using SPSS 26 statistical software (SPSS Inc, Chicago, IL, USA).

## Accession numbers

Sequence information from this article can be found in the Arabidopsis Genome Initiative or GenBank/EMBL databases under the following accession numbers: SWEET5 (AT5G62850), GALK (AT3G06580), USP (AT5G52560).

## Acknowledgments

We thank Dr. Yaxin Li for help with sample collection and thank Dr. Qi Xie for providing us with p*YAO:hSpCas9* vector. We also thank Mrs. Anita K. Snyder for editing. This work is supported by startup funds from the University of Illinois at Urbana-Champaign to Dr. Chen (to J.W., Y.-C. Y., and L.-Q.C). Y. L. was supported by China Scholarship Council (No. 201603250070).

